# Stem cell therapy for skin regeneration using mesenchymal stem cells derived from the progeroid Werner syndrome-specific iPS cells

**DOI:** 10.1101/2021.06.15.448474

**Authors:** Shinichiro Funayama, Hisaya Kato, Hiyori Kaneko, Kentaro Kosaka, Daisuke Sawada, Aki Takada-Watanabe, Takuya Minamizuka, Yusuke Baba, Masaya Koshizaka, Akira Shimamoto, Yasuo Ouchi, Atsushi Iwama, Yusuke Endo, Naoya Takayama, Koji Eto, Yoshiro Maezawa, Koutaro Yokote

**Affiliations:** Department of Endocrinology, Hematology and Gerontology, Chiba University Graduate School of Medicine, Chiba, Japan; Division of Diabetes, Metabolism and Endocrinology, Chiba University Hospital, Chiba, Japan; Department of Regenerative Medicine, Chiba University Graduate School of Medicine, Chiba, Japan; Department of Plastic, Reconstructive, and Aesthetic Surgery, Chiba University Graduate School of Medicine, Chiba, Japan; Department of Pediatrics, Chiba University Graduate School of Medicine, Chiba, Japan; Department of Medicine, Division of Diabetes, Endocrinology and Metabolism, Kimitsu Chuo Hospital, Kisarazu, Chiba, Japan; Department of Regenerative Medicine Research, Faculty of Pharmaceutical Sciences, Sanyo-Onoda City University, Sanyo-Onoda, Yamaguchi, Japan; Gene Expression Laboratory, Salk Institute for Biological Studies, La Jolla, CA, USA; Division of Stem Cell and Molecular Medicine, Center for Stem Cell Biology and Regenerative Medicine, The Institute of Medical Science, The University of Tokyo, Tokyo, Japan; Laboratory of Medical Omics Research, KAZUSA DNA Research Institute, Kisarazu, Chiba, Japan; Department of Clinical Application, Center for iPS Cell Research and Application (CiRA), Kyoto University, Kyoto, Japan

**Keywords:** Werner syndrome, wound healing, mesenchymal stem cell, iPS cell, aging

## Abstract

Adult progeria, Werner syndrome (WS), is an autosomal recessive disorder that develops accelerated aging-associated symptoms after puberty. Refractory skin ulcer of limbs, which is one of the symptoms specific to WS, is seriously painful and sometimes results in amputation. In recent years, cell therapy using mesenchymal stem cells (MSCs) has been attracting attention; however, the effect of WS-derived MSCs on skin ulcers is still unclear. In this study, we generated iPS cells from a patient with WS and a normal subject, differentiated them into MSCs (WS- and NM-iMSC, respectively), and performed cell therapy to a refractory skin ulcer mouse model. As a result, WS-iMSC recapitulated premature senescence phenotypes in vitro. Upon subcutaneous injection around the wounds of mice, WS-iMSC was significantly inferior in wound healing effect compared to NM-iMSC. Proteome and transcriptome analysis revealed altered expression of genes related to angiogenesis, inflammation, and proliferation in WS-iMSC with remarkable downregulation of VEGF, a potent angiogenic factor. In addition, simultaneous administration of recombinant human VEGF and WS-iMSC improved the wound healing effect in vivo. These results indicate that the expression of angiogenic factors is reduced in WS-iMSC, and its supplementation restores the wound healing ability. This finding may pave the way to develop the treatment of intractable skin ulcers of WS.

## Introduction

Werner syndrome (WS), caused by mutation of a RecQ type helicase gene *WRN*, is an autosomal recessive progeroid syndrome that causes various signs of accelerated aging after puberty, including bilateral cataracts, graying and loss of hair, type 2 diabetes, sarcopenia, dyslipidemia, arteriosclerosis, and malignant tumors [1, 2]. In addition to these symptoms, WS patients exhibit disease-specific phenotypes such as calcification of the Achilles tendon, refractory skin ulcers, and high susceptibility to non-epithelial tumors, e.g., sarcomas and hematological malignancies [3, 4]. Among those, refractory painful skin ulcers occur in about 70% of WS patients and often lead to amputation of the lower limbs accompanied by severe pain and osteomyelitis, which result in remarkably reduced patient quality of life [5]. Owing to the lack of a fundamental cure for this condition, the development of a treatment strategy is urgently needed.

Mesenchymal stem cells (MSCs) are somatic stem cells that possess the ability to differentiate into mesenchymal lineages, such as osteocytes, chondrocytes, and adipocytes [6–8]. MSCs have been clinically applied for a wide range of diseases and reported to be effective for graft-versus-host disease, stroke, multiple sclerosis, and diabetic skin ulcers [9–12]. Therefore, the clinical application of MSCs is considered a promising tool for regenerative medicine [13]. On the other hand, MSCs derived from aged mice reportedly had reduced wound healing effects compared to those derived from young mice [14]. Related to this issue, previous reports suggested MSCs derived from human *WRN* knock-out embryonic stem cells exhibited premature senescence phenotypes [15]. Taken together, although treatment with MSCs may have beneficial effects for refractory skin ulcers in WS patients, MSCs derived from WS patients are considered to have decreased wound healing effects than those derived from normal individuals because of accelerated senescence. However, the effect of MSCs from WS on ulcer treatment remains to be elucidated.

To clarify the skin regenerative effect of MSCs from WS, we generated iPS cells from a normal subject and a patient with WS, differentiated them into MSCs (NM- and WS-iMSC, respectively), and performed cell transplantation using a refractory skin ulcer mouse model.

## Results

### Generation of NM- and WS-iPSC

iPSCs were generated from a normal individual and a patient with WS as previously described [16]. They showed iPSC-like morphologies and the capability of embryoid body formation (Supplementary Figure 1A, B). Gene expression analysis revealed the downregulation of pluripotency genes and upregulation of genes of three germ layers upon differentiation (Supplementary Figure 1C). Sanger-sequencing showed compound heterozygous *WRN* mutation, c.3139-1G>C plus c.3446delA, in WS-iPSC (Supplementary Figure 1D). Thus, we successfully generated NM- and WS-iPSC.

### WS-iMSC recapitulated premature senescence phenotypes in vitro

Derivation of iMSCs was conducted according to the previous report with modification [17]. Derived iMSCs exhibited spindle-shaped MSC-like morphologies (Supplementary Figure 2A). They were positive for CD73, CD90, and CD105 and possessed adipogenic, chondrogenic, and osteogenic potentials (Supplementary Figure 2B, C). During the long-term culture, WS-iMSC showed decreased proliferative capacity compared to NM-iMSC (Figure 1A). The analysis of telomere length displayed shortened telomeres in WS-iMSC (Figure 1B). SA-β-gal analysis disclosed a significantly increased number of senescent cells in WS-iMSC (Figure 1C, D). These results indicate that WS-iMSC recapitulated premature senescence phenotypes in vitro.

**Figure 1.**
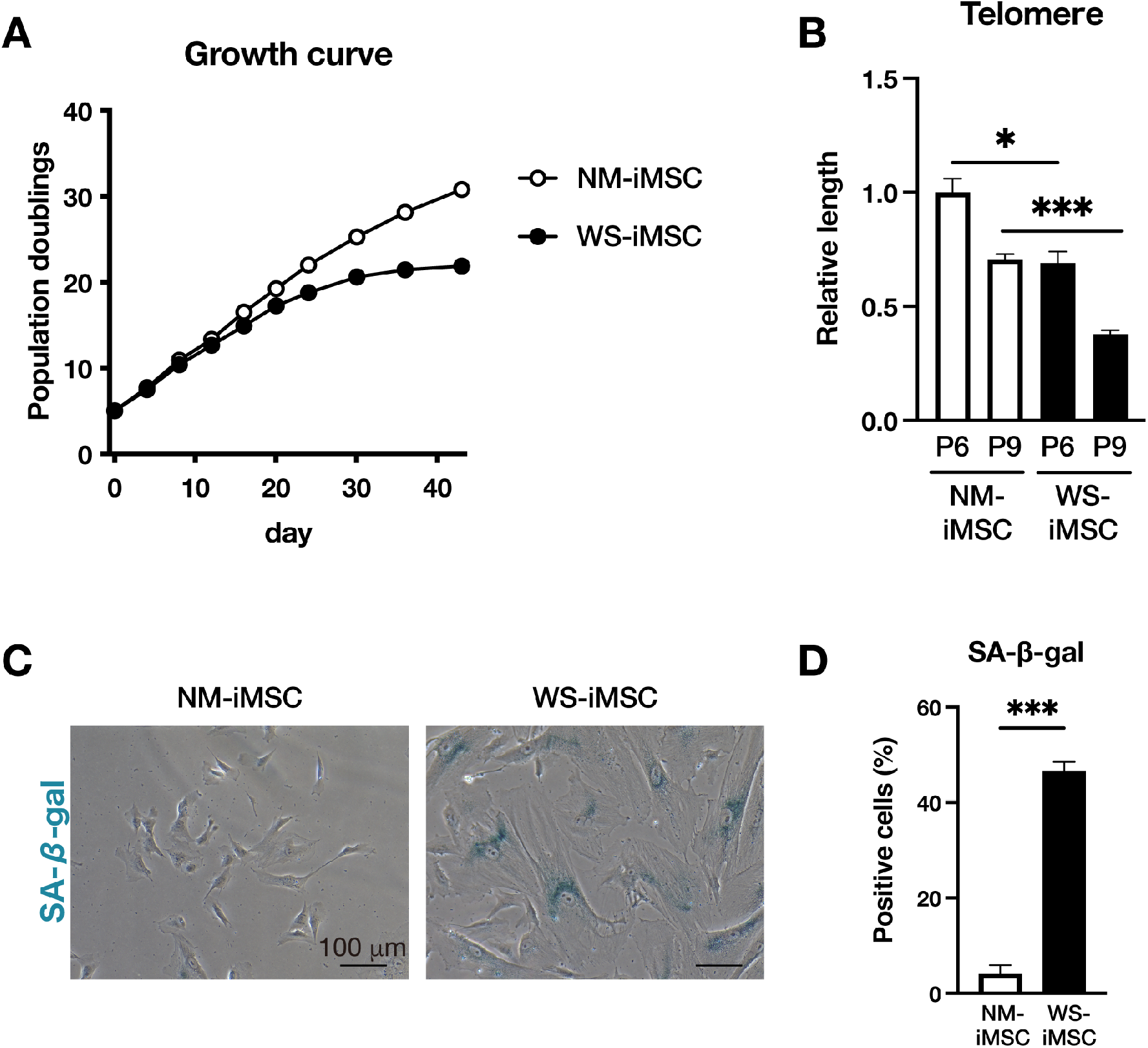
Premature senescence phenotypes of WS-iMSC. (A) Growth curves of NM- and WS-iMSC. (B) Relative telomere length quantified by qPCR at passages 6 and 9. Data are mean ± SEM of three technical replicates. A student t-test was performed (*p<0.05, ***p<0.001). (C) Representative images of SA-β-gal staining at passage 12. Scale bar = 100 μm. (D) The rate of positive cells related to Figure 1C. More than 200 cells in each group were counted. Data are mean ± SEM of five microscopic views. A student t-test was performed (***p<0.001).

### WS-iMSC exhibited reduced wound healing effects compared to NM-iMSC

Refractory skin ulcer mouse models were generated by administering streptozotocin (STZ) to severe combined immunodeficient (SCID) mice and creating a wound on their back, as previously reported (Figure 2A) [18]. To determine the wound healing effects of iMSCs, NM- and WS-iMSC suspended in hyaluronic acid were subcutaneously injected into the area around the wound, and wound sizes were tracked for 14 days. As a result, mice administered NM-iMSC showed significantly decreased wound size compared to the groups of vehicle and WS-iMSC on day 14 (Figure 2B, C). Consistent with this finding, attenuated epidermal and dermal thickness was observed in mice administered WS-iMSC (Figure 2D, E, F, G). These results suggest that WS-iMSC possesses insufficient wound healing effects compared to NM-iMSC.

**Figure 2.**
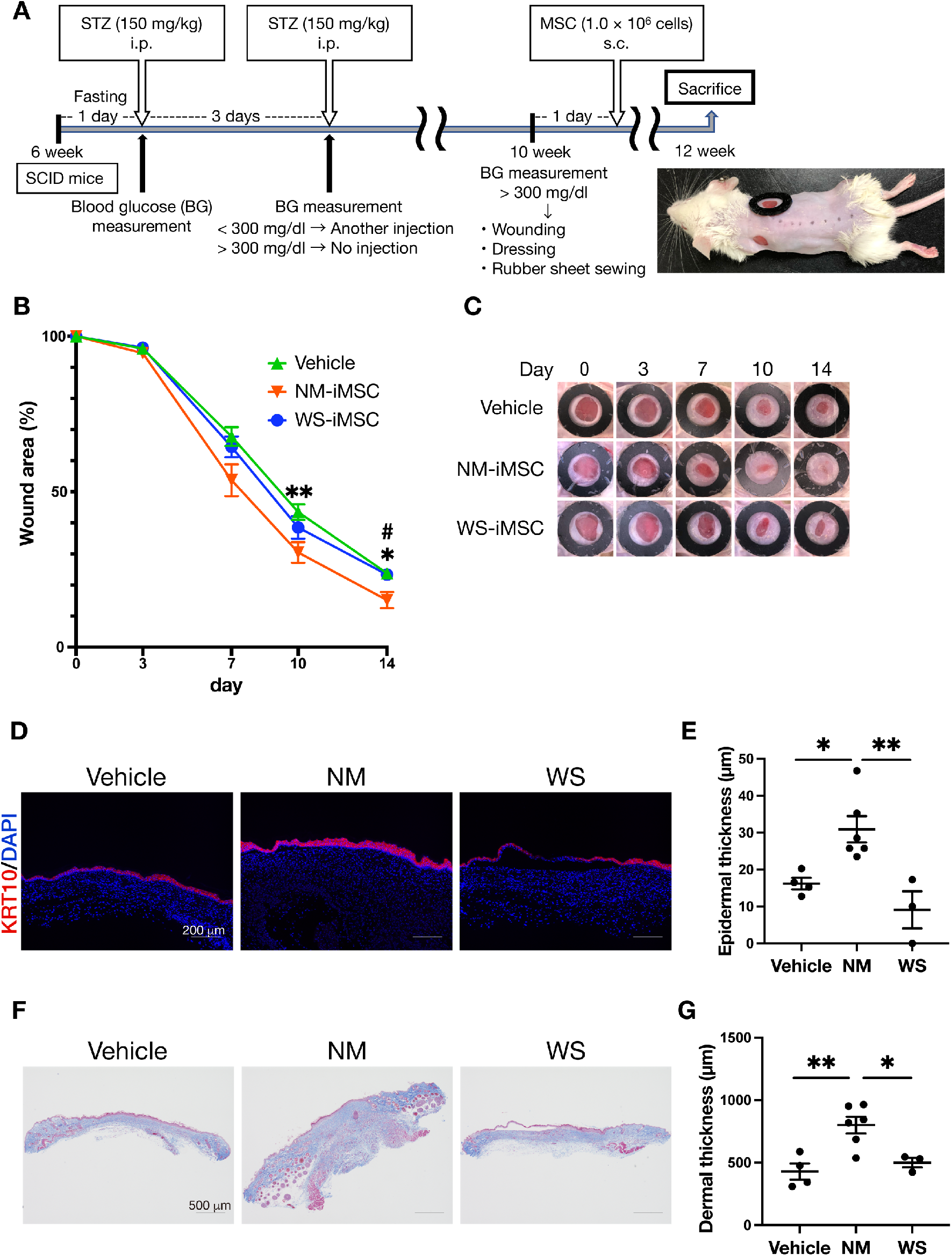
Wound healing effects of refractory skin ulcer mouse model by administering NM- and WS-iMSC. (A) Scheme representing the production of refractory skin ulcer mouse model and subsequent experiments. (B) Graph showing the rate of wound area from day 0 to 14. Data are mean ± SEM (Veh, n = 8; NM, n = 10; WS, n = 6). A student t-test was performed (*p<0.05, **p<0.01, Veh vs NM-iMSC; #p<0.05, WS-iMSC vs NM-iMSC). (C) Representative pictures of wounds on mice in each group. (D) Representative images of immunohistochemical staining with KRT10 and DAPI of mouse skin sections on day 14. (E) Quantification of epidermal thickness related to Figure 2D. Data are mean ± SEM. A student t-test was performed (*p<0.05, **p<0.01). (F) Representative images of mouse skin sections stained by Masson’s trichrome staining method. (G) Quantification of dermal thickness related to Figure 2F. Data are mean ± SEM. A student t-test was performed (*p<0.05, **p<0.01).

### NM-iMSC facilitated angiogenesis

To assess the angiogenic effects of iMSCs, distributions of mouse Pecam-1 and Vegf expression in dermal skin sections on day 14 were evaluated using the in situ hybridization method. The dermis of mice administered NM-iMSC showed increased and decreased expression of Pecam-1 and Vegf, respectively (Figure 3A, B, C). On the other hand, mice administered WS-iMSC and vehicle exhibited opposite outcomes, meaningly expressions of decreased Pecam-1 and increased Vegf in the dermis (Figure 3A, B, C). Conversely, in the analysis of epidermis, expression of Vegf was elevated in mice administered NM-iMSC and reduced in those of WS-iMSC and vehicle (Figure 3D, E). These findings suggest that the angiogenesis was facilitated by NM-iMSC, while the distribution of Vegf expression in the skin after the administration of WS-iMSC is distinct from that of NM-iMSC.

**Figure 3.**
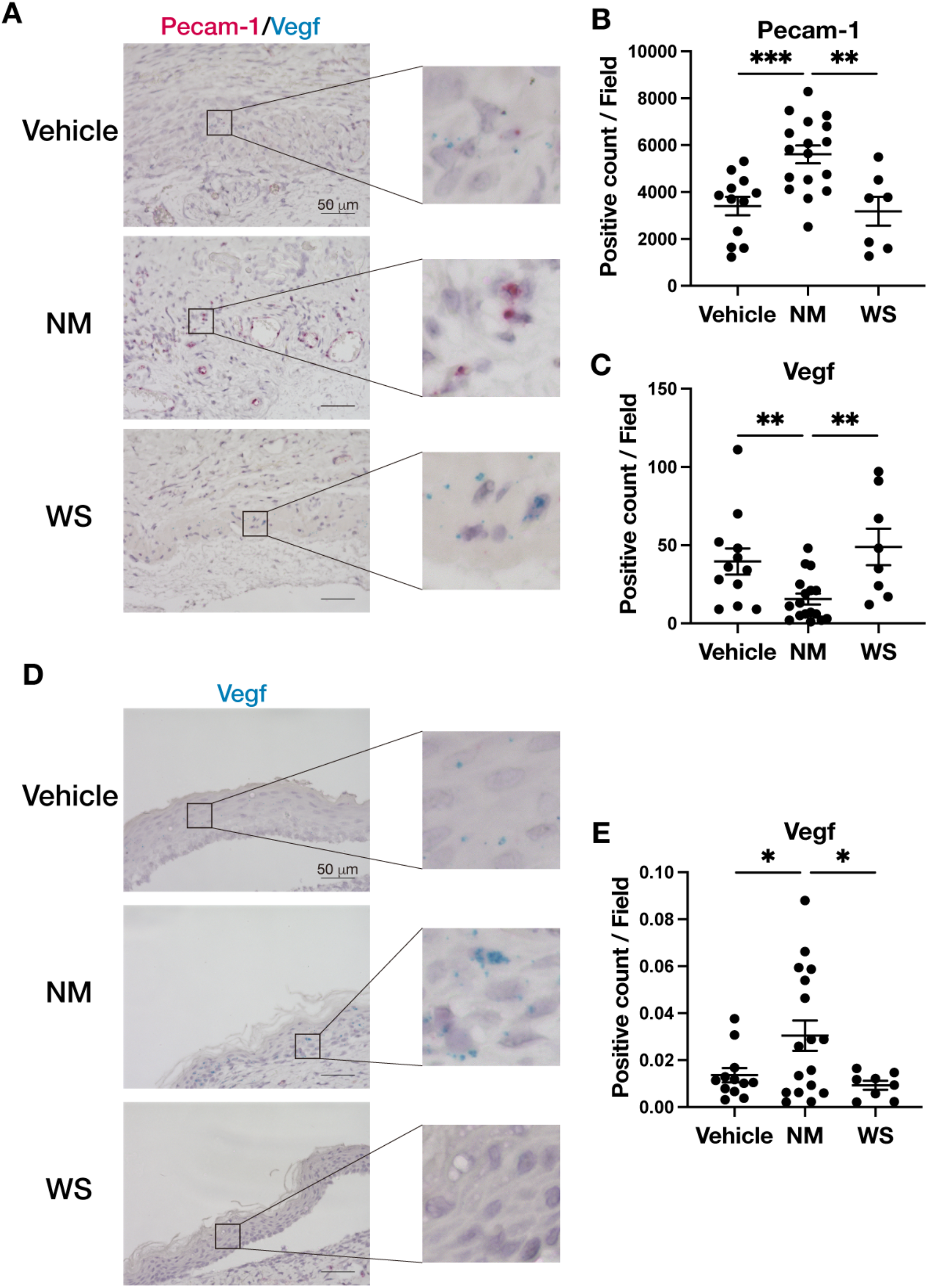
Analysis of Pecam-1 and Vegf expression in dermis and epidermis by in situ hybridization. (A) Representative images of Pecam-1 (red dots) and Vegf (blue dots) in dermal sections of mice on day 14. (B), (C) Quantification of dots positive for Pecam-1 (B) and Vegf (C) related to Figure 3A. Data are mean ± SEM. A student t-test was performed (**p<0.01, ***p<0.001). (D) Representative images Vegf (blue dots) in epidermal sections of mice on day 14. (E) Quantification of dots positive for Vegf related to Figure 3D. Data are mean ± SEM. A student t-test was performed (*p<0.05).

### NM-iMSC promoted the migratory effect of WS-fibroblasts compared to WS-iMSC in vitro

Fibroblasts are responsible for producing extracellular matrix to promote angiogenesis [19]. Further, fibroblasts have pivotal roles in the wound healing process at the cell proliferation stage [20]. Rapid migration of fibroblasts to the wound site and their proliferation are major components in the acceleration of wound healing [21–23]. Thus, we conducted a co-culture experiment to determine whether co-culturing fibroblasts with NM- or WS-iMSC can promote their migration ability in vitro using Transwell. As a result, there was no difference in the migration ability between dermal fibroblasts from a normal subject (NM-fibroblasts) co-cultured with NM-iMSC and those co-cultured with WS-iMSC (Figure 4A, B). However, the migration ability of fibroblasts from a WS patient (WS-fibroblasts) was significantly reduced when co-cultured with WS-iMSC compared to NM-iMSC (Figure 4A, B). These results suggest that WS-iMSC is inferior to NM-iMSC in its ability to secrete factors that promote the migration of WS-fibroblasts.

**Figure 4.**
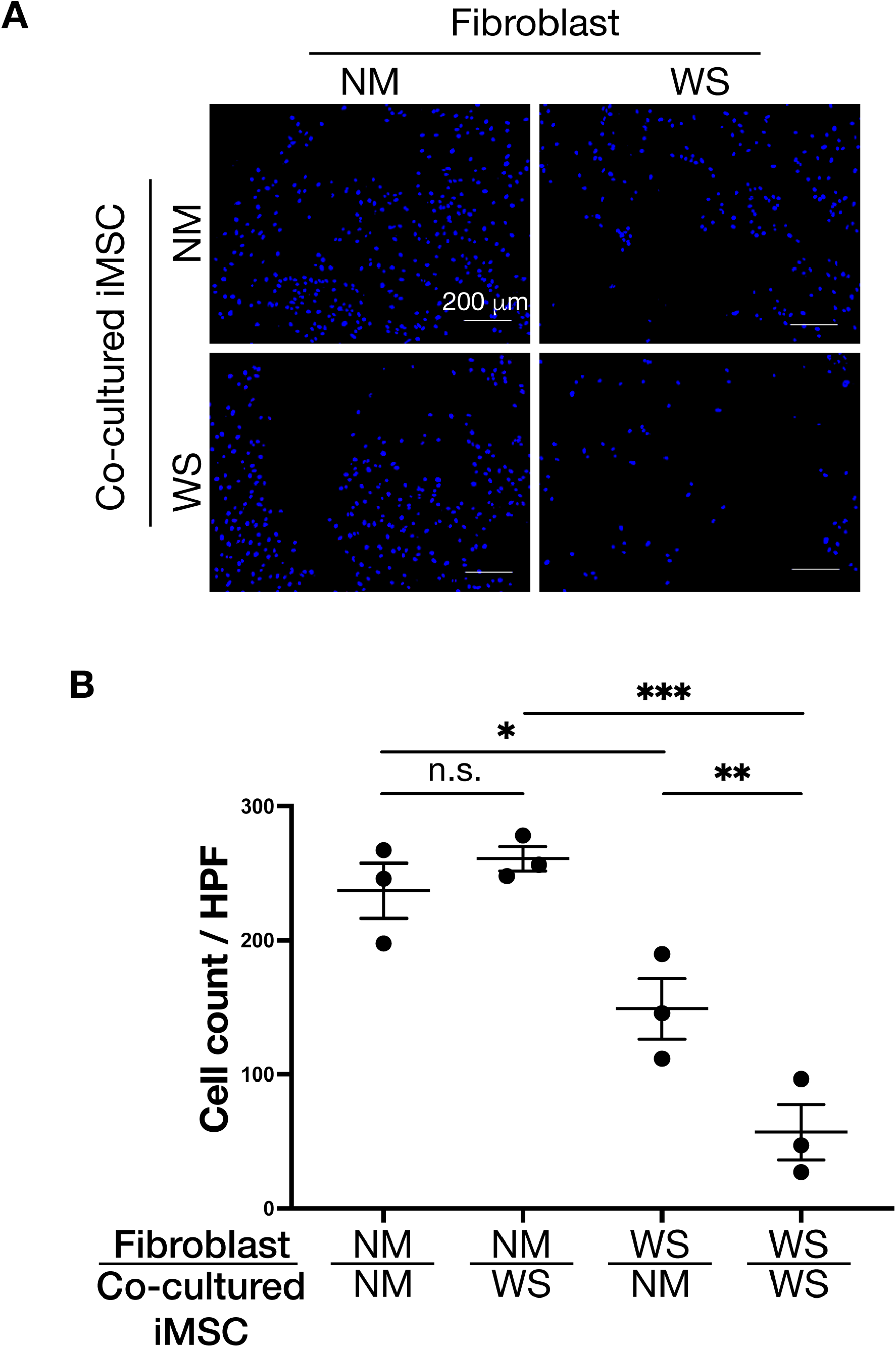
Analysis of the migratory ability of fibroblasts co-cultured with NM- and WS-iMSC. (A) Representative images of fibroblasts that migrated beneath the Transwell chamber stained with DAPI after 24-hour co-culturing with NM- and WS-iMSC. (B) Quantification of migrated fibroblasts related to Figure 4A. Data are mean ± SEM (n = 3). A student t-test was performed (n.s., not significant; *p<0.05; **p<0.01; ***p<0.001).

### NM- and WS-iMSC showed distinct secretome regarding wound healing

MSCs are known to promote angiogenesis and cell proliferation by secreting various cytokines and chemokines [24, 25]. Based on the above results of co-culture experiments, the secreted factors from iMSCs were assumed to be associated with fibroblasts migration and wound healing. Therefore, we carried out a proteome analysis of the culture supernatant of iMSCs to clarify their wound healing-associated secretome. As a result, WS-iMSC had significantly increased secretion of the inflammation-related factors; IL-8, MCP-1, Pentraxin3, and MMP-9 compared to NM-iMSC (Figure 5A, B, C). On the other hand, the secretion of proteins related to cell proliferation, such as Activin A, IGFBP-3, and VEGF, were significantly upregulated in NM-iMSC (Figure 5A, B, C). Indeed, enzyme-linked immunosorbent assay confirmed significantly lower VEGF secretion in WS-iMSC than in NM-iMSC (Figure 5D). These findings suggest that NM-iMSC and WS-iMSC have distinct secretomes associated with wound healing.

**Figure 5.**
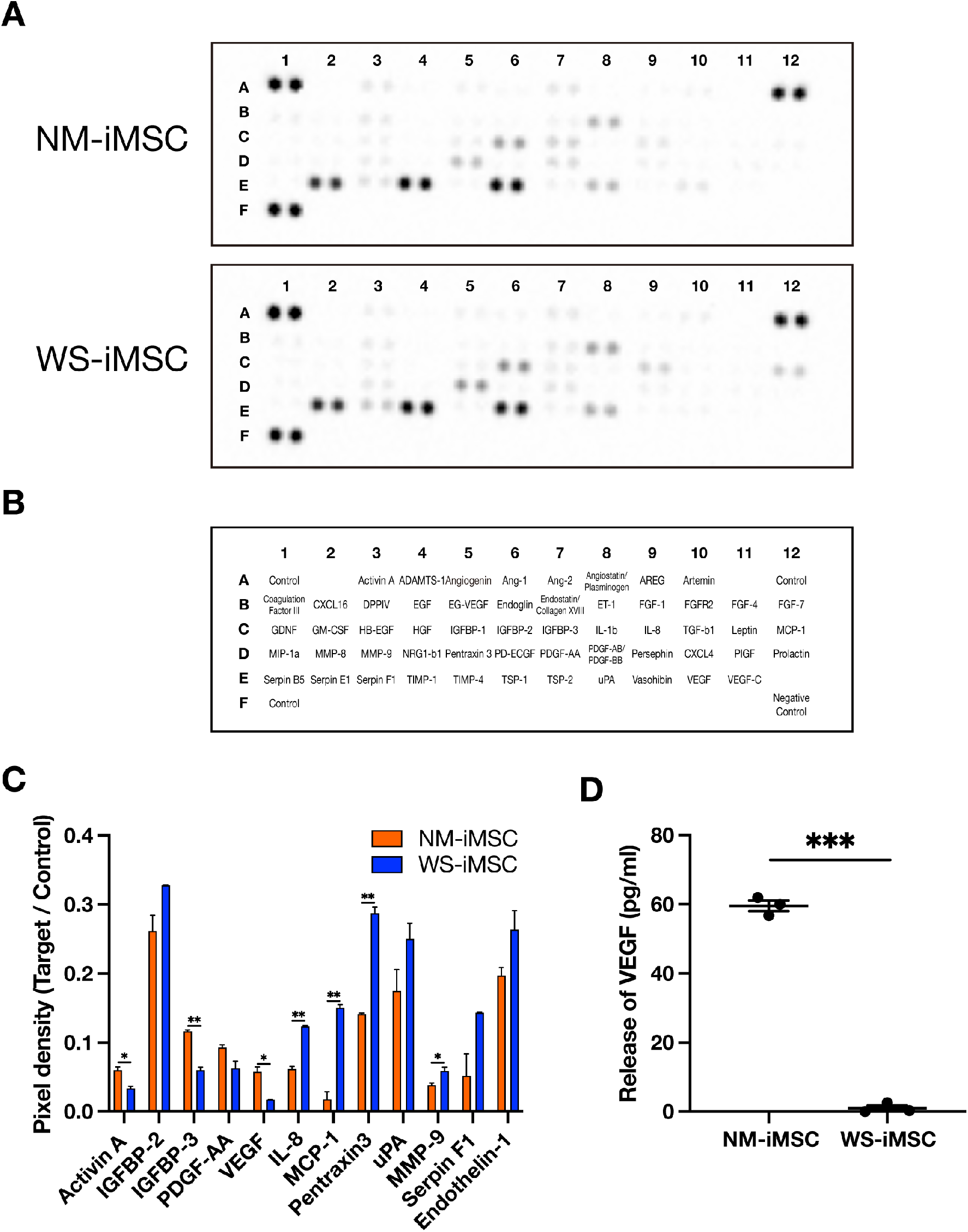
Analysis of secreted factors in NM- and WS-iMSC conditioned medium. (A) Images of the blotted membrane in proteome analysis of angiogenic factors secreted by NM- and WS-iMSC. (B) Plot table of the membrane in Figure 5A. (C) Quantification of proteome analysis shown in Figure 5A. Data are mean ± SEM. A student t-test was performed (*p<0.05; **p<0.01). (D) The concentration of VEGF in conditioned medium measured by ELISA. Data are mean ± SEM. A student t-test was performed (***p<0.001).

### Gene expression profiles related to angiogenesis and inflammation were altered in WS-iMSC

Next, we evaluated the transcriptome of NM- and WS-iMSC via RNA-sequence. As a consequence, we extracted 1,114 and 886 genes that were respectively downregulated or upregulated in WS-iMSC compared to NM-iMSC (Figure 6A). Enrichment analysis revealed genes involved in chromosome segregation and cell division were downregulated in WS-iMSC, while those of anatomical structure morphogenesis and regulation of multicellular organismal process were upregulated (Figure 6B). Especially, the expression levels of genes associated with angiogenesis and inflammation, such as VEGFA, CXCL8, etc., were remarkably altered (Supplementary Table). qRT-PCR confirmed significantly different gene expression profiles of the above genes between NM- and WS-iMSC (Figure 6C). These findings might be associated with the attenuated regenerative capacity of the refractory skin ulcer in WS-iMSC.

**Figure 6.**
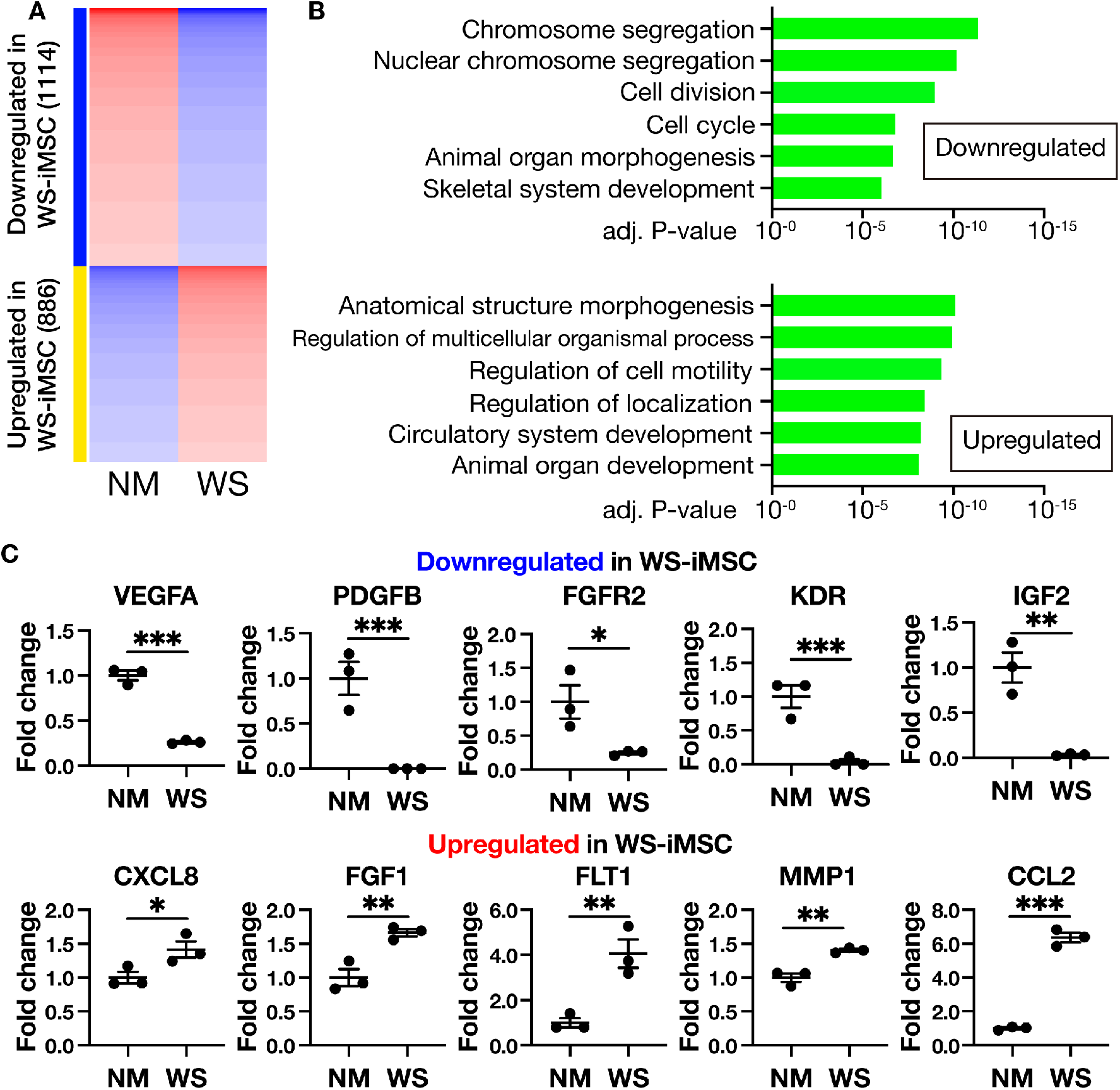
Gene expression analysis of NM- and WS-iMSC. (A) Heatmap of differentially expressed genes between NM- and WS-iMSC in transcriptome analysis. (B) List of gene ontology (GO: biological process) analysis and corresponding p-values related to Figure 6A. (C) qRT-PCR results of genes related to cell proliferation, angiogenesis, and inflammation. Data are means ± SEM of three technical replicates. A student t-test was performed (*p<0.05; **p<0.01; ***p<0.001).

### VEGF supplementation improved the wound healing effect of WS-iMSC

VEGF is a potent angiogenic factor and is also essential in wound healing [26, 27]. Since VEGF expression was reduced in WS-iMSC, we speculated that VEGF supplementation might improve the wound healing effect of WS-iMSC. Therefore, WS-iMSC and recombinant human VEGF were co-injected around the wound of the refractory skin ulcer mouse model. As a result, the wound area on day 10 was significantly reduced in the VEGF injected group (Figure 7A, B) compared to WS-iMSC alone. From these results, the decreased wound healing ability of WS-iMSC might attribute to the deficiency of VEGF secretion.

**Figure 7.**
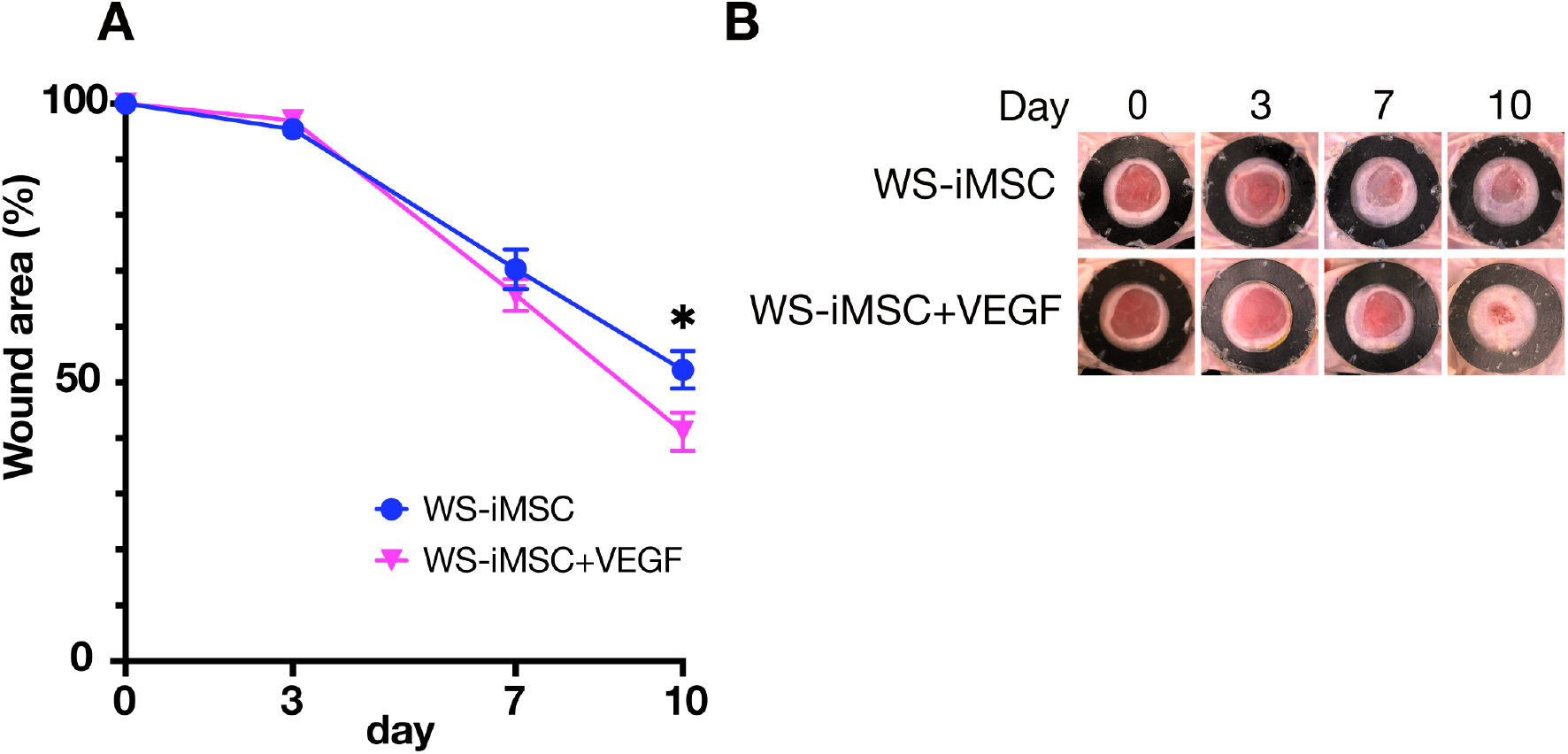
Wound healing effects of administration of WS-iMSC alone or WS-iMSC plus VEGF on intractable skin ulcers. (A) Graph showing the rate of wound area from day 0 to 10. Data are mean ± SEM (n = 7). A student t-test was performed (*p<0.05). (B) Representative pictures of wounds on mice in each group.

## Discussion

In this study, we demonstrated the difference in wound healing efficacy between NM- and WS-iMSC when administered into a refractory skin ulcer mouse model. WS-iMSC manifested premature senescence phenotypes in vitro and weaker wound healing effects compared to NM-iMSC resulting in thinner epidermal and dermal layers in injected mice. Further, mice administered WS-iMSC had decreased and increased expression of mouse Pecam-1 and Vegf in the dermis, respectively, and, in contrast, decreased Vegf expression in the epidermis compared to those administered NM-iMSC. WS-fibroblasts co-cultured with WS-iMSC exhibited reduced migration ability in vitro compared to those co-cultured with NM-iMSC. The proteome analysis revealed significantly reduced VEGF secretion in WS-iMSC. The transcriptome analysis also confirmed significantly lower expression of genes involved in cell proliferation, blood vessels, and inflammation, including VEGFA, in WS-iMSC. Indeed, VEGF supplementation restored the wound healing effect of WS-iMSC.

There are four phases in wound healing: hemostasis, inflammation, proliferation, and remodeling, and various chemokines and cytokines are secreted in each phase [28]. Among those, it is well established that VEGF plays one of the most important roles in angiogenesis [29]. Previous reports suggest that VEGF secreted by myeloid cells and fibroblasts is highly expressed in the dermis in the early stages of wound healing to promote angiogenesis, but its expression shifts to the epidermis in the late stage as keratinocytes migrate onto granulation tissue [28–30]. Therefore, the observed difference in our study in the expression distribution of Vegf in skin sections of mice administered NM- and WS-iMSC might indicate the delayed progression of the wound healing phase in mice administered WS-iMSC.

The relationship between VEGF and the therapeutic effects of MSCs has been demonstrated in several previous studies. Concomitant injection of MSCs and VEGF reduced the infarct size and preserved ejection fraction in a mouse model of acute myocardial infarction [31]. MSCs overexpressing VEGF demonstrated to promote the migration of vascular endothelial cells and increase vascular density and blood flow in hindlimb ischemic mice [32]. Of note, VEGF proved to be effective in prolonging the MSC survival at the injected site [33]. These findings not only indicate that VEGF secreted by MSCs plays an important role in promoting angiogenesis but also suggest a protective effect of VEGF for MSCs. In the present study, RNA sequencing and proteome analysis revealed significantly lower VEGF expression in WS-iMSC than in NM-iMSC, and simultaneous injection of VEGF and WS-iMSC improved wound healing ability. Therefore, the decreased secretion of VEGF in WS-iMSC might have contributed to the delay in wound healing, and its restoration might ameliorate the regenerative ability of WS-iMSC.

Few reports have described the relationship between VEGF and WS so far. Goto M, et al. reported an increased serum VEGF level in WS patients [34]. Another study indicated that knockdown of the WRN gene under a hypoxic condition led to an increased VEGF expression due to HIF1α augmentation in HeLa cells [35]. On the other hand, MSCs collected from older people have reportedly decreased VEGF secretion [36]. WS-iMSC showed reduced VEGF secretion in our study, but further studies are needed to elucidate the underlying mechanism.

In this study, WS-iMSC exhibited reduced treatment capacity to refractory skin ulcers, probably due to the decreased secretion of VEGF. These findings might contribute to the elucidation of disease pathogenesis and the development of novel therapeutic approaches in the near future.

## Materials and Methods

### iPSC and iMSC induction and cell culture

Cell culture was performed at 37 °C with 5% CO_2_ under humidified air. iPSCs were generated from a male normal individual and a male patient with WS in their 50s as previously described [16]. Derivation of iMSCs was conducted according to the previous report with modification [17]. Briefly, embryoid bodies were cultured in collagen type IV-coated plates (Corning, 354416) with MSC derivation medium (alpha-MEM (Invitrogen), 10% FBS (Hyclone), Antibiotic-Antimycotic (Gibco, 15240062), 100 nM dexamethasone (Sigma-Aldrich), 50 uM L-ascorbic acid 2-phosphate sesquimagnesium salt hydrate (Sigma-Aldrich, A8960-5G)). After five days, cells were passaged on collagen type I (Nitta Gelatin, Cellmatrix Type I-C) fibril-coated plate (defined as passage 0, population doubling level 0) and cultured in expansion medium (alpha-MEM (Invitrogen), 10% FBS (Hyclone), Antibiotic-Antimycotic (Gibco, 15240062), non-essential amino acids (Invitrogen, 11140-050)). The medium change was performed every 3 days. When reaching subconfluent, cells were passaged at a 1:4 split ratio. To draw the growth curve, 5 × 10^4^ of NM- and WS-iMSC at PDL5 (P3) were seeded on collagen I coated 6 well plates (Iwaki, 4810-010). 5 × 10^4^ cells were counted and passaged every 4 to 5 days until P12. Dermal fibroblasts, established from a male normal individual or a male WS patient in their 40s, were cultured in DMEM (043-30085, Wako) supplemented with 10% FBS (10270106, Gibco) and antibiotic (15240062, Gibco) in normal culture dishes (TR4002, TrueLine) at 37 °C with 5% CO_2_. The cells at subconfluency were passaged at a ratio of 1:4, and those with population doublings (PD) of 8 to 18 were used in the experiment.

### Fluorescence-activated cell sorting (FACS)

Surface markers of MSCs were analyzed by flow cytometer (BD FACSCanto II). In detail, MSCs were dispersed using Trypsin-EDTA (Gibco, 25200072) and suspended in PBS with 2mM EDTA and 0.5% BSA (Sigma, A3294-50G). Cells were incubated with primary antibodies (anti-CD73 (BD Biosciences, 550257), anti-CD90 (BD Biosciences, 555595), and anti-CD105 (eBioscience, 17-1057-42)) for 30 min at room temperature and analyzed.

### Tri-lineage differentiation

In vitro differentiation potentials of MSCs into three lineages were evaluated by using adipogenesis, chondrogenesis, and osteogenesis differentiation kit (A1007001, A1007101, and A1007201, respectively. All from Gibco) according to the manufacturer’s protocols. For each assay, oil red o, alcian blue, and alizarin red stainings (All Sigma) were used, respectively.

### Relative telomere length measurement

For the analysis of the relative telomere length, genomic qPCR was conducted using the SYBR Green PCR master mix (Applied Biosystems), as previously described [37].

### Senescence-associated β-galactosidase staining

Senescence-associated β-galactosidase (SA-β-Gal) staining was performed following the manufacturer’s instruction (Cell Signaling, 9860S). After staining the nuclear DNA using Hoechst 33,342 (DOJINDO, 346–07951), the positive rate was calculated.

### Breeding environment

Mice were housed under a temperature of 24 ± 2°C and humidity of 55 ± 5%, with light exposure from 6:00-18:00 (12-hour automatic lighting). Aspen chips were placed in the plastic cage as bedding, and mice were fed CE-2 (Oriental Yeast Co. Ltd.), a solid feed for mice.

### Preparation of refractory skin ulcer model mice and administration of iMSC

Diabetes mellitus was induced in six-week-old severe combined immunodeficient (SCID) mice (C.B17/Icr-scidJcl scid/scid, CLEA Japan, Inc.), as previously described [18]. Briefly, mice were administered 150 mg/kg of streptozotocin (STZ; S0130-1G, Sigma-Aldrich). At three days after STZ administration, the blood glucose (BG) was measured, and mice with BG ≧ 300 mg/dl were considered diabetic immunodeficient mice (DM-SCID). Mice that did not reach the BG of 300 mg/dl received a second administration of STZ (150 mg/kg), and their BG was measured three days later.

Ten-week-old DM-SCID mice were anesthetized, and a 6-mm wound was created on the skin using the Disposable Biopsy Punch (BP-60F, Kai Medical). A donut-shaped rubber sheet with internal and external diameters of 10 mm and 16 mm, respectively, created by cutting out a 1-mm thick silicone rubber sheet (Kyowa Industries), was sewn around the wound of mice. Perme-roll (H24R05, Nitto Medical) was applied for wound dressing.

Three experimental groups of mice were established: vehicle group, normal group, and WS group. For the vehicle group, 600 *μ*l of sodium hyaluronate (6.66 mg/ml, H0603, Tokyo Chemical Industry) was mixed with 100 *μ*l of DMEM (043-30085, Wako Pure Chemical Industries). Thereafter, a total of 100 *μ*l of the mixture was subcutaneously injected around the wound divided into four points. For the NM and WS groups, 7.0×10^6^ NM- or WS-iMSC suspended in 100 *μ*l of DMEM was further suspended in 600 *μ*l of hyaluronic acid. Thereafter, a total of 100 *μ*l of cell suspension (including 1.0×10^6^ cells) was injected around the wound divided into four points. The wound was photographed on days 0, 3, 7, 10, and 14 after the injection, and mice were dissected on day 14. The wound areas were analyzed using the image analysis software, AreaQ. In the VEGF co-administration experiment, 0.15 ng of human recombinant VEGF (Gibco) were co-injected with 1.0×10^6^ cells of WS-iMSC.

### Immunohistochemical staining

The skin around the wound was isolated with a disposable biopsy punch (6 mm) at the time of dissection. The isolated skin was then cut into two pieces, one of which was fixed with 4% paraformaldehyde to prepare a paraffin-embedded section (GenoStaff). After deparaffinization and rehydration, the prepared slides were immersed in 10 mM citric acid buffer, autoclaved for antigen activation, and immersed in 3% H2O2/PBS to inhibit the endogenous peroxidase activity. After blocking, anti-Cytokeratin 10 antibodies (ab76318, Abcam) and Alexa Flour 594 (1:1000, A11037, Thermo Fisher) were added to the sections. The sections were sealed with DAPI-containing Fluoro-KEEPER (12745-74, Nacalai Tesque) and analyzed using the image analysis software ImageJ.

### HE staining and Masson’s trichrome staining

Paraffin-embedded sections were deparaffinized and rehydrated. Thereafter, the sections were stained with Mayer’s hematoxylin solution (032-14635, Wako). After staining with eosin solution, the sections were lyophilized and sealed. Masson’s trichrome staining was performed by GenoStaff. The sections were analyzed using the image analysis software ImageJ.

### In situ hybridization

In situ hybridization was performed using the RNAscope Duplex Color Assay Kit (RNAscope^(R)^2.5 HD Duplex Reagent Kit, 322430, ACD), according to the product protocol. After baking at 60 °C in a HybEZ oven (HybEZ™ Hybridization System With EZ-Batch Slide System, 321461, ACD), the slides were deparaffinized and rehydrated. After treated with H2O2, the slides were immersed in RNAscope^(R)^ Target Retrieval Reagent (322000, ACD) heated to 100-104 °C for 15 min for antigen activation. Protease treatment of the sections by using Protease Plus was performed at 40 °C for 30 min. The mixture probe, prepared by mixing RNAscope® Probe-Mm-Vegfa-ver2 (412261, ACD) and RNAscope® Probe-Mm-Pecam1-C2 (316721-C2, ACD) at a ratio of 50:1, was added to the sections for hybridization at 40 °C for 2 h. The signal was further amplified by incubation. The Pecam-1 and VEGF signals were stained with red and blue, respectively. The sections were analyzed using the image analysis software ImageJ.

### Transwell migration assay

NM- and WS-iMSC were seeded at a density of 6.0×10^4^ cells/well in 24-well plates. After culturing for 24 h, a culture insert (for 24-well plates, 8.0 *μ*m, 353097, Corning) was attached to each well, and 2.0×104 NM- or WS-fibroblasts were seeded on the insert. After 24 h, cells on top of the insert were removed with a cotton swab, and cells migrated into the bottom of the insert were fixed with 4% paraformaldehyde. The membrane of the insert was removed and attached to a slide, and finally sealed with a DAPI-containing Fluoro-KEEPER. The analysis was performed using the image analysis software ImageJ.

### Proteome analysis and Enzyme-linked immunosorbent assay (ELISA)

NM- and WS-iMSC were trypsinized and seeded at a density of 1.5×10^5^ cells/well in 12-well plates. After culturing for 24 h, the medium was replaced with 1 ml of MEMα, and cells were cultured for another 24 h. Thereafter, the culture supernatant was collected and stored at −80 °C. For analysis, the Proteome Profiler Human Angiogenesis Array Kit (511-61901, Wako) was used according to the protocol. Blotted membrane was analyzed using ChemiDocTMMP (BIO-RAD) and Image Lab. By using the same samples, we carried out an ELISA, according to the protocol of the VEGF Human ELISA Kit (ab100662, Abcam).

### Gene expression analysis

A total of 100,000 NM- and WS-iMSC collected in 1.5-ml tubes were pelleted for storage at −80 °C. RNA was extracted from the cells by using TRIzol^(R)^Reagent (15596026, ambion), according to the protocol of the PureLink™ RNA Micro Scale Kit (12183016, Thermo Fisher). RNA sequencing was performed by the Kazusa DNA Research Institute. The obtained FASTQ file was mapped to the human genome, GRCh38.99, using STAR (ver.2.7.6a) and RSEM (ver.1.3.3) to obtain the BAM file. The obtained gene count data were analyzed using iDEP [38]. Gene clustering was performed by analyzing the top 2,000 genes with variable expression by using k-Means. For qRT-PCR, cDNA was synthesized as previously described [39]. Following probes were used (all from TaqMan); VEGFA, Hs00900055_m1; PDGFB, Hs00966522_m1; FGFR2, Hs01552926_m1; KDR, Hs00911700_m1; IGF2, Hs01005963_m1; CXCL8, Hs00174103_m1; FGF1, Hs01092738_m1; FLT1, Hs01052961_m1; MMP1, Hs00899658_m1; CCL2, Hs00234140_m1; GAPDH, Hs02786624_g1 (internal control).

### Study approval

All experiments were approved by the institutional review boards at the Chiba University Graduate School of Medicine (Chiba, Japan). Written informed consent was obtained from study participants before the commencement of this research.

## Author Contributions

S.F. and H.Kato designed the study, carried out the experiments, analyzed the data, wrote the manuscript, and composed the figures; H.Kaneko, K.K., D.S., A.T.W., T.M., and Y.B., carried out the experiments; M.K., A.S., Y.O., A.I., N.T., and K.E. discussed the data; Y.E. conducted transcriptome analysis; Y.M. and K.Y. designed the study, discussed the data, and managed funding; all authors approved the final version of the manuscript.

## Acknowledgments

This work was supported by Japan Society for the Promotion of Science (JSPS) KAKENHI under Grant Numbers JP19K23939 (H.Kato), JP20H00524 (K.Y.), JP20K16542 (H.Kato); Japan Agency for Medical Research and Development (AMED) under Grant Numbers JP20bm0804016 (K.Y.), JP20ek0109353 (K.Y.), JP20gm5010002 (K.Y.); Ministry of Health, Labour and Welfare (MHLW) of Japan under Grant Number H30-nanchi-ippan-009 (K.Y.).

---

**Supplementary Figure 1.**
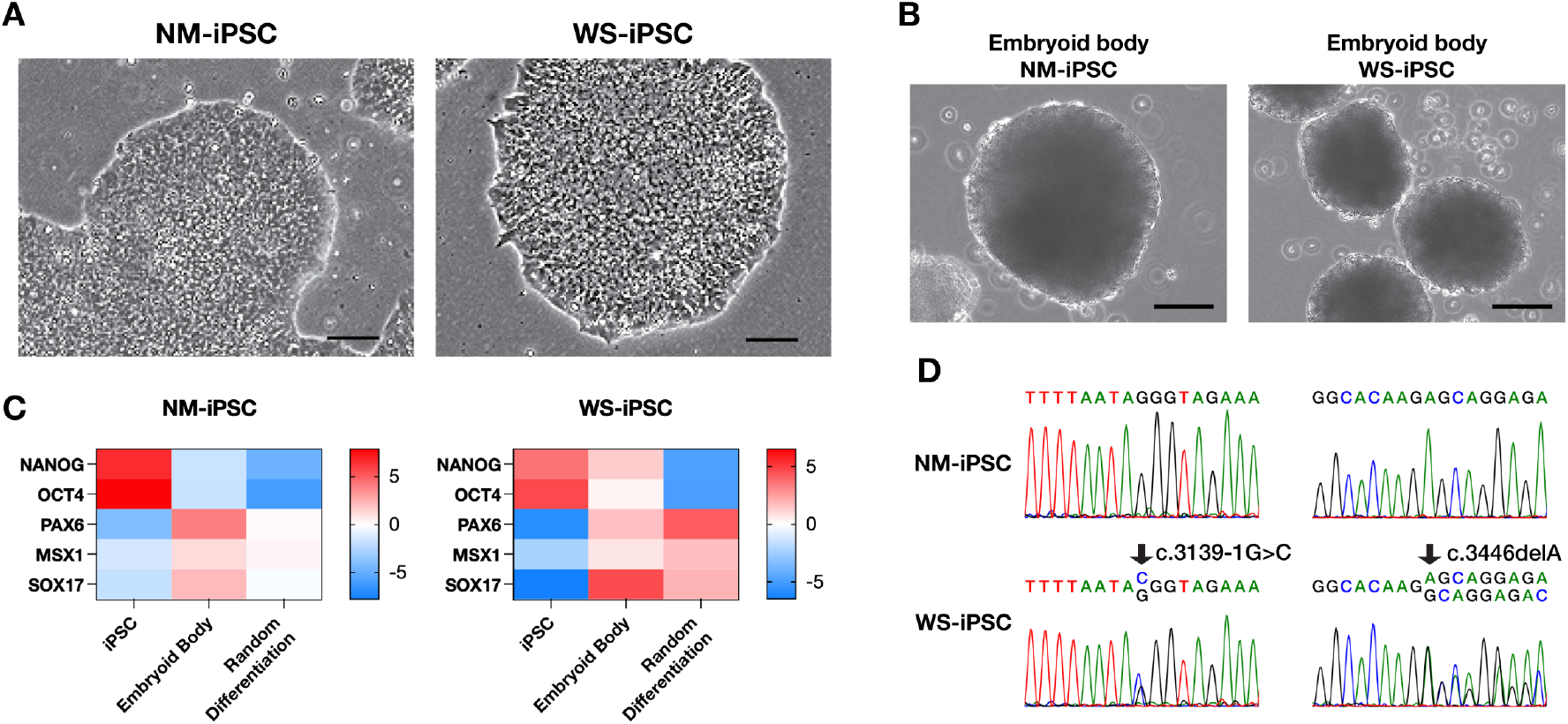
Characteristics of NM- and WS-iPSC. (A) Representative images of iPSCs. Bar = 100*μ*m. (B) Representative images of embryoid bodies. Bar = 100*μ*m. (C) Heatmap of qRT-PCR results assessing genes of pluripotency and three germ layers. (D) Sanger sequencing results at *WRN* mutated locus.

**Supplementary Figure 2.**
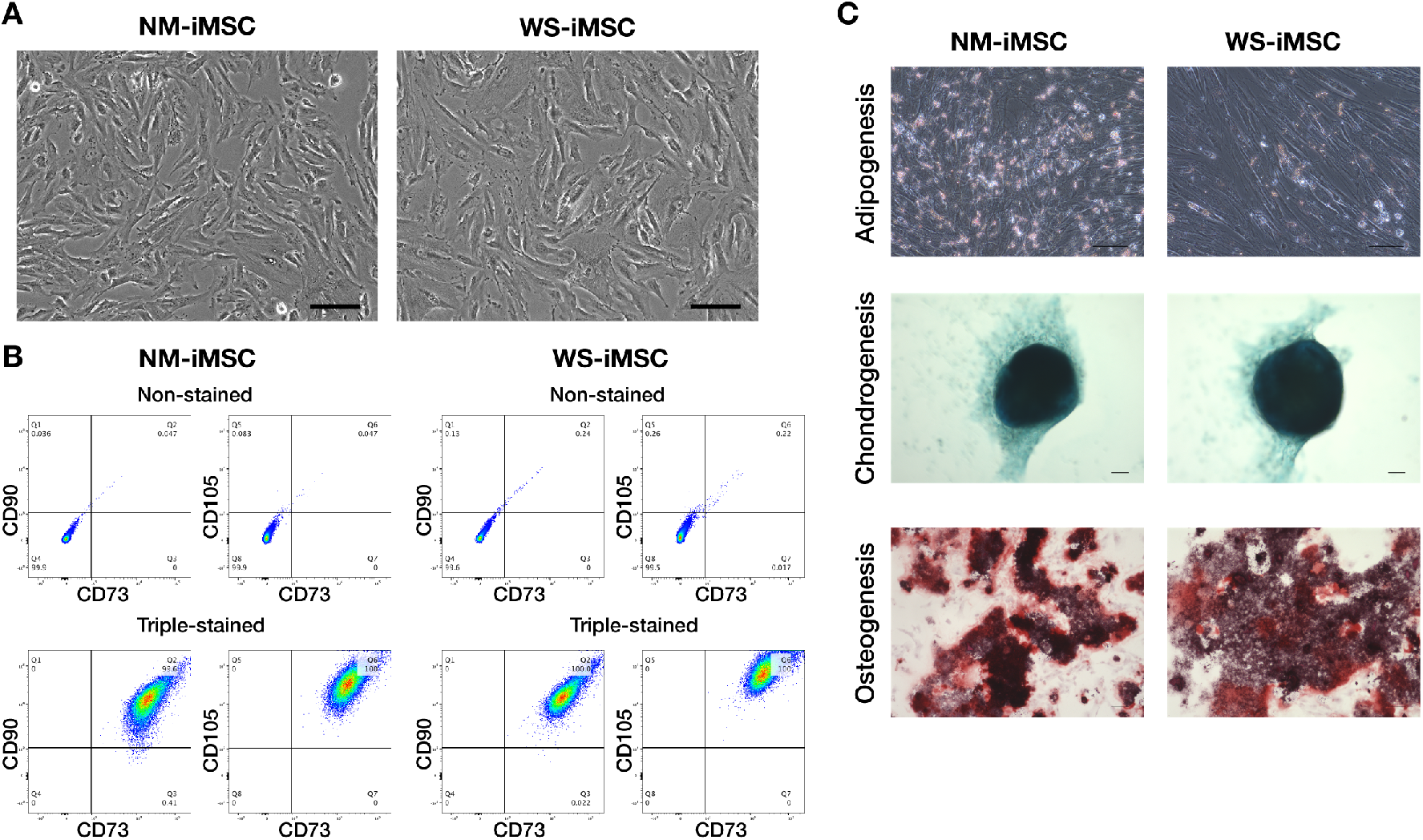
Characteristics of NM- and WS-iMSC. (A) Representative images of iMSCs at PD8. Bar = 100 *μ*m. (B) FACS quantification of positive cell rates for cell surface markers specific to MSCs. (C) Representative images of tri-lineage differentiated cells. For staining, oil red O to adipogenesis, alcian blue to chondrogenesis, and alizarin red to osteogenesis were used. Bar = 100 *μ*m.

Supplementary Table. A list of the top 2000 genes with the highest SD in RNA-sequence of NM- and WS-iMSC.

